# Striatal cholinergic and dopaminergic driven astrocyte Ca^2+^ activity is disrupted in Parkinsonian mice

**DOI:** 10.64898/2026.07.08.737296

**Authors:** Wesley R. Evans, Hunter G. Wells, Cynthia Jacob, Angelica Vellore, Rafiq Huda

## Abstract

Brain neuromodulatory systems exert powerful effects on local neuronal circuit function and behavior. In addition to classical actions directly on neurons, growing evidence indicates that neuromodulators also recruit Ca^2+^-dependent astrocyte mechanisms to regulate synaptic plasticity and network function. The dorsal striatum integrates cortical and thalamic inputs under strong dopamine (DA) and acetylcholine (ACh) neuromodulatory control. To what extent the circuit and behavioral effects of striatal ACh and DA depend on astrocyte Ca^2+^ activity remains unclear. We show that locomotion elicits robust DA, ACh, and astrocyte Ca^2+^ activity in the dorsolateral striatum (DLS). DA and ACh release exhibits a negative correlation on a fast time scale but shows a positive correlation during continuous locomotion as slower astrocyte Ca^2+^ activity builds. Higher ACh and DA release is associated with higher astrocyte events, suggesting that both neurotransmitters drive astrocyte activity. In agreement, pharmacological blockade of muscarinic ACh or D1/D2 DA receptors decreases locomotion-evoked astrocyte Ca^2+^. Closed-loop optogenetic inhibition of striatal cholinergic interneurons (CINs) during locomotion reduces astrocyte Ca^2+^ activity, demonstrating a causal contribution of ACh release to astrocyte activity. Locomotion related ACh release was severely compromised in a mouse model of Parkinson’s disease (PD), with the dual loss of DA and ACh attenuating astrocyte Ca^2+^ activity. Facilitating astrocyte cholinergic signaling via chemogenetics improved both calcium activity and motor deficits in our recent work. Thus, the pathophysiology of PD in part involves attenuated astrocyte Ca^2+^ signaling, placing these non-neuronal cells as a prime underexplored therapeutic target for PD.

## Introduction

Neuromodulation in the CNS is a critical component of neural circuit function and exerts wide-ranging effects depending on context and neurotransmitter identity^1–4^. Across brain regions, neuromodulators such as norepinephrine (NE), DA, and ACh have effects on memory formation, behavior, and circuit activty^5–7^ Astrocytes, a non-neuronal glial cell common across all regions of the CNS, have emerged in recent years as a local mediator of neuromodulatory activity^8–16^. Astrocytes exhibit robust Ca^2+^ activity which reflects the activity of local neurons and neurotransmitter release^17–19^. Neuropathological conditions change astrocyte genetic profiles and Ca^2+^ activity, demonstrating that these cells respond to pathological conditions^20–24^.

The striatum is the input nucleus of the basal ganglia that integrates thalamic, cortical, and substantia nigra pars compacta (SNpc) DA activity and is involved in motor, habitual, and other behaviors^5,25,26^. In particular, the DLS is a site of convergence for primary motor cortex and thalamic inputs involved in the execution of motor behavior^25^. Loss of striatal DA innervation during PD leads to imbalanced activity of striatal projection neurons (SPNs)^5,27–29^. The striatum also contains its own population of CINs which release ACh in constant fluctuation with striatal DA activity^30,31^. Together, these two neuromodulatory systems form an important signaling axis in the striatum which is impaired when DA levels drop such as in PD^30–35^. The striatum is largely devoid of locus coeruleus NE projections and activity^36,37^, leaving the ACh-DA axis as a major candidate for communication to astrocytes given the established NE-astrocyte mechanisms in other brain regions^38–42^. However, the relationship between DA, ACh, and astrocyte Ca^2+^ in the DLS is yet to be fully elucidated.

We recently showed^23^ reduced astrocyte Ca^2+^ activity in a mouse model of PD (unilateral DA denervation with 6-OHDA). We investigated the upstream elements of striatal circuitry that drive astrocyte Ca^2+^ rises and which may explain the suppression of astrocyte Ca^2+^ we observed. We began our investigation with the hypothesis that the striatal astrocytes are attuned to either ACh or DA, assuming only one of these systems is the primary communication axis to astrocytes. Instead, we found that locomotion elicits ACh and DA release along with astrocyte Ca^2+^ activity. ACh and DA rises both are associated with increased astrocyte Ca^2+^. We further show that pharmacological blockade of muscarinic and dopaminergic receptors both lower locomotion-evoked astrocyte calcium and that optogenetic silencing of CINs acutely lowers astrocyte calcium levels. Finally, we demonstrate that DA denervation with 6-OHDA leads to dysfunctional locomotion-related ACh release and astrocyte Ca^2+^. Counteracting this ACh deficit by directly augmenting astrocyte cholinergic signaling led to improvements in both astrocyte Ca^2+^ activity and PD-relevant motor behavior in our recent work^23^. Together, these results show the functional relevance of astrocyte Ca^2+^ signaling for PD motor deficits and provide further evidence of the therapeutic potential of these non-neuronal cells.

## Results

### 1. DA, ACh, and astrocyte Ca^2+^ activities display a consistent temporal pattern during locomotion

We employed a two-site fiber photometry recording strategy to measure ACh or DA release together with astrocyte Ca^2+^ activity. We virally expressed in one DLS hemisphere GCaMP6f-cyto in astrocytes and either GRAB-DA2m or GRAB-ACh3.0 in the contralateral hemisphere (**Fig 1a**). Bilateral astrocyte activity was found to be highly correlated (**Fig. S1**). DA and ACh release also shows a high bilateral correlation^31^, allowing us to analyze contralateral astrocyte GCaMP activity as an approximation of in-site simultaneous activity. After surgical recovery, mice were trained to voluntarily locomote on a circular wheel while head-fixed (**Fig 1b**). GRAB and GCaMP6f activity were simultaneously recorded alongside locomotor data allowing us to detect onsets of voluntary movements and assess the aligned ΔF/F signal (**Fig 1c-e**). Around locomotion onset, we detected an ACh peak alongside a fast DA dip, similar to that observed during transitions in spontaneous behavior^43^. These initial changes were followed by a sustained increase in both ACh and DA release for the duration of the locomotion bout, a period during which astrocyte Ca^2+^ activity slowly rose (**Fig 1f, g**). After locomotion offset, ACh and DA levels both fall, with astrocyte activity beginning to decline soon after (**Fig 1f**). To better isolate the DA response associated with continuous locomotion after the initial dip, we performed a baseline correction by subtracting activity just before locomotion onset (**Fig 1h**). This showed that after dipping, DA activity largely mirrors the resultant rise in astrocyte Ca^2+^ (**Fig 1i**). These data evidence a stereotyped course of events after locomotion onset in which ACh and DA show opposed activity on a fast timescale but then remain at elevated levels for the duration of locomotion. This period while both neuromodulators are elevated coincides with the rising phase of astrocyte Ca^2+^. Thus, both ACh and DA activity exhibit relationships to subsequent astrocyte Ca^2+^ increases, necessitating we investigate the possibility that the two neuromodulatory systems may work in tandem.

**Fig 1:**
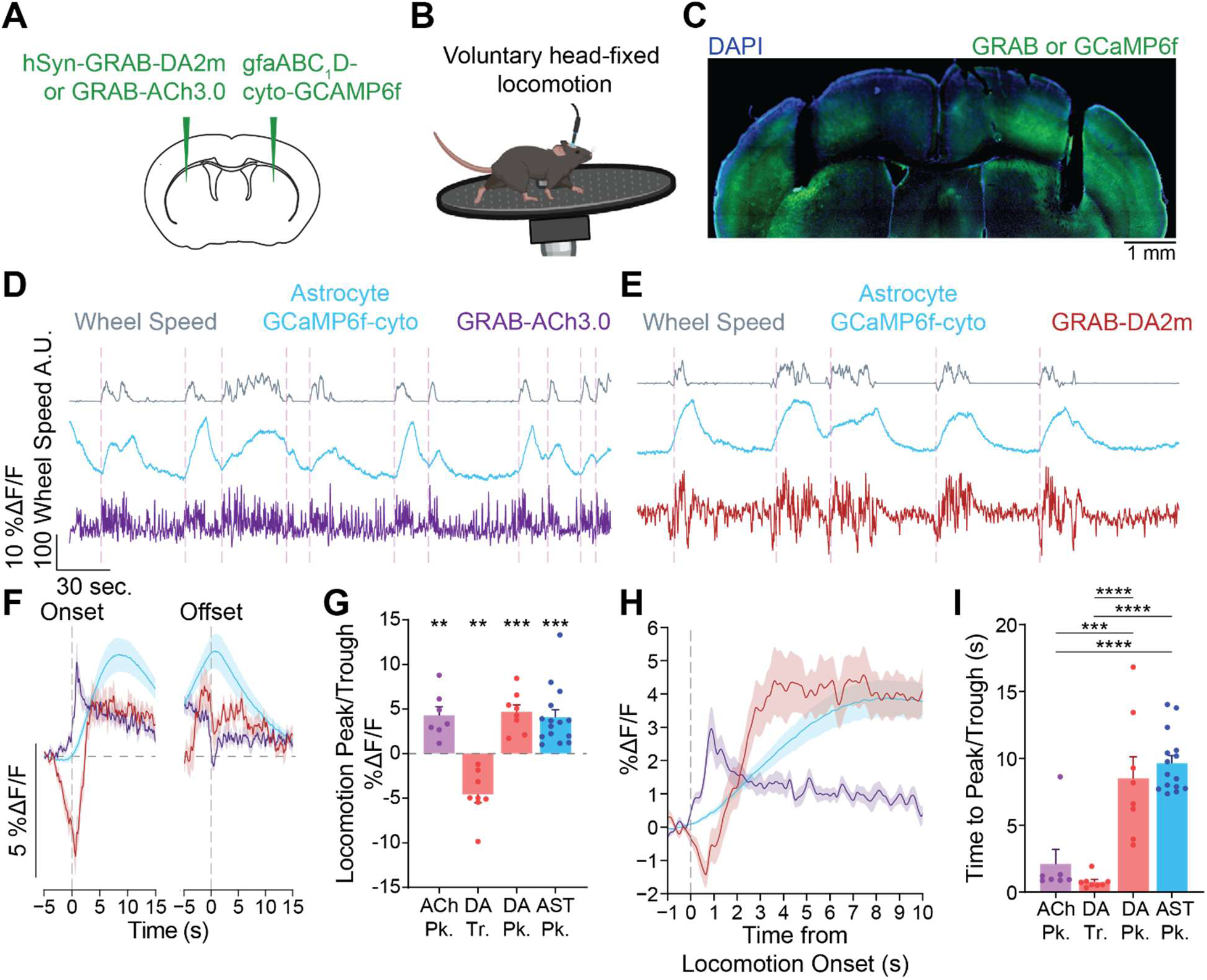
Striatal DA, ACh, and astrocyte Ca^2+^ activity all exhibit locomotion-induced tonic shifts. (**A**) Experimental schematic showing bilateral viral expression strategy for co-recording of astrocyte Ca^2+^ with gfaABC_1_D-cyto-GCaMP6f contralateral to either hSyn-GRAB-DA2m or hSyn-GRAB-ACh3.0. (**B**) Schematic of head-fixed voluntary wheel apparatus for measurement of locomotor activity. (**C**) Epifluorescence tile-scan image of bilateral fiber placement and viral expression in DLS. (**D**) Example recording showing five minutes of simultaneous locomotor, astrocyte Ca^2+^, and ACh activity with detected locomotion onsets indicated by dashed lines. (**E**) The same as (**D**) showing 5 minutes of co-recorded locomotor, astrocyte Ca^2+^, and DA activity. (**F**) Animal averaged ACh, DA, and astrocyte Ca^2+^ ΔF/F aligned to locomotion onset (left) and offset (right). Vertical dashed line represents locomotion onset, horizontal represents baseline 0 value. (**G**) Magnitude of detected ACh peaks (Pk.), DA peaks and troughs (Tr.), and astrocyte Ca^2+^ (AST) peaks (One-sample t-test against 0: ACh peak value *p* = 0.0044, DA trough value *p* = 0.0019, DA peak value *p* = 0.00059, astrocyte Ca^2+^ peak *p* = 0.00028, n = 7 GRAB-ACh3.0 mice, 8 GRAB-DA2m mice, and 15 astrocyte GCaMP6f mice). (**H**) animal averaged ACh, DA, and astrocyte Ca^2+^ ΔF/F baseline corrected by the mean of activity from -1s to 0s showing locomotion-induced activity shifts. (**I**) Time to reach either indicated signal peak or trough relative to locomotion onset (One-way ANOVA *F*(3, 34) = [24.00], *p* = 1.6 × 10^−8^; Tukey’s multiple comparisons test: time to ACh peak vs. DA trough *p* = 0.79, ACh peak vs DA peak *p* = 0.00059, ACh peak vs AST peak *p* = 8.2 × 10^−6^, DA trough vs DA peak *p =* 2.3 × 10^−5^, DA trough vs. AST peak *p* = 1.6 × 10^−7^, DA peak to AST peak *p* = 0.80).

### 2. Simultaneous recordings of astrocyte Ca^2+^, DA, and ACh reveals ACh and DA release predicts subsequent astrocyte activity

Our initial experiments reveal that both ACh and DA are potential mediators of astrocyte Ca^2+^ activity. To test this possibility, we simultaneously recorded ACh, DA, and astrocyte Ca^2+^ levels with fiber photometry by virally introducing rDA1m and rACh1.7 into opposite striatal hemispheres of transgenic mice expressing GCaMP8s in astrocytes (**Fig 2a**). This strategy led to robust overlap of construct expression (**Fig 2b**). We validated GCaMP8s expression by counterstaining striatal tissue for the astrocyte marker S100β and the neuronal marker NeuN (**Fig 2c**). This experimental strategy allowed us to record the status of ACh, DA, and astrocyte Ca^2+^ simultaneously in the same animal (**Fig 2d**). We again utilized a voluntary locomotor paradigm to align fiber photometry signals to locomotion onset and offset to assess inter-signal correlation<u>s</u> (**Fig 2e, f**). Trial-averaged ACh and DA release was negatively correlated around locomotion onset but showed a net positive correlation for the remainder of the locomotion bout. We also observed that DA and astrocyte Ca^2+^ activity were relatively correlated after locomotion onset, concurring with our observations in **Fig 1i**. ACh and astrocyte activity showed a transient increase in correlation around locomotion onset after which their activity was negatively correlated. To test how ACh and DA relate to subsequent astrocyte Ca^2+^ activity, we separated individual locomotion trials into quartiles based on relevant metrics of ACh and DA activity. When stratified by trial ACh peak ΔF/F, resulting astrocyte ΔF/F peaks were larger in the highest quartile than in the lowest quartile (**Fig 2g** left, **h**). When stratified by either DA dip depth or peaks, astrocyte ΔF/F peaks were only larger in the highest vs. lowest quartiles by peaks (**Fig 2g** center, **i**) and not when separated by dip magnitude (**Fig 2g** right, **j**). On single trials, peak ACh and DA ΔF/F were moderately correlated with astrocyte Ca^2+^ responses (**Fig 2k, l**). Astrocyte activity showed a weak negative correlation with the magnitude of DA dips around locomotion onset (**Fig 2m**). Our observations strengthen support for a model where both DA and ACh increases recruit astrocyte Ca^2+^ activity, albeit at different time points post-locomotion. An initial increase in ACh release seems to trigger the astrocyte Ca^2+^ response which slowly builds during sustained release of both ACh and DA. Conversely, DA dip depth appears less predictive of subsequent astrocyte Ca^2+^ activity. Thus, increased levels of both DA and ACh are associated with increased subsequent astrocyte Ca^2+^ events.

**Figure 2:**
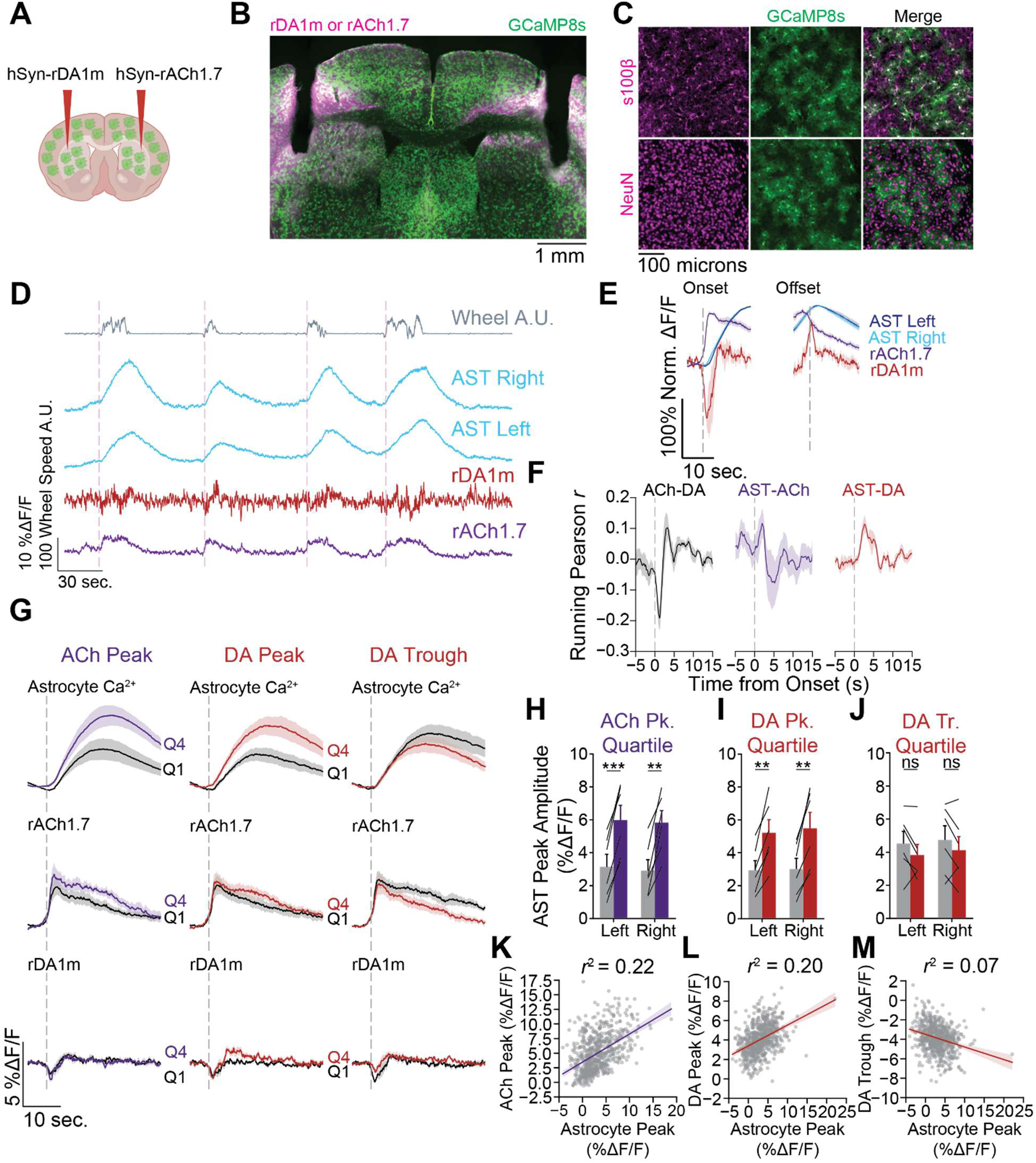
Multicolor fiber photometry reveals correlation of astrocyte Ca^2+^ activity to increases in striatal ACh and DA levels. (**A**) Schematic of viral construct injection strategy to bilaterally express hSyn-rDA1m and hSyn-rACh1.7 in mice transgenically expressing GCaMP8s in astrocytes. (**B**) Epifluorescence tile-scan image of recording fiber placement in DLS with expression of astrocyte GCaMP8s and either hSyn-rDA1m or hSyn-rACh1.7. (**C**) Expression of astrocyte GCaMP8s counterstained against the astrocyte maker S100β and the neuronal marker NeuN. (**D**) Example trace showing five minutes of simultaneous recording of bilateral astrocyte Ca^2+^ and unilateral DA and ACh activity with detected locomotion onsets indicated by dashed lines. (**E**) ACh, DA, and astrocyte Ca^2+^ activity expressed as ΔF/F normalized to session averaged trial max aligned to locomotion onset and offset. (**F**) Running 2s window of Pearsons *r* value visualizing locomotion-induced correlational epochs of ACh and DA (left), ACh and astrocyte Ca^2+^ (center), and DA and astrocyte Ca^2+^ (right). (**G**) Stratification of trials by ACh peak ΔF/F (left), DA peak ΔF/F (center), and DA trough ΔF/F (right) displaying the locomotion onset aligned animal averaged traces of the highest (Q4) and lowest (Q1) quartiles for astrocyte Ca^2+^ (top), ACh (middle), and DA (bottom). (**H**) Quantification of astrocyte Ca^2+^ peak ΔF/F from trials in either the highest (Q4) or lowest (Q1) ACh peak quartiles (left hemisphere Q1 vs Q4 paired t-test *p* = 0.00019, right hemisphere *p* = 0.0016, n = 6 mice). (**I**) Same as (**H**) with quartiles of DA peak ΔF/F (left hemisphere Q1 vs. Q4 paired t-test *p* = 0.0022, right hemisphere *p* = 0.0019, n = 6 mice). (**J**) Same as (**H**) with quartiles of DA trough (left hemisphere Q1 vs. Q4 paired t-test *p* = 0.75, right hemisphere *p* = 0.57, n = 6 mice). (**K**) Scatterplot and best fit line of all trials ACh peak ΔF/F and astrocyte Ca^2+^ peak ΔF/F, Pearson’s *r*^2^ = 0.22, *p* = 1.6 × 10^−37^ n = 663 trials. (**L**) Same as (**K**) with DA peak ΔF/F and astrocyte Ca^2+^ peak ΔF/F, Pearson’s *r*^2^ = 0.20, *p* = 2.2 × 10^−27^. (**M**) Same as (**K**) with DA trough ΔF/F and astrocyte Ca^2+^ peak ΔF/F, Pearson’s *r*^2^ = 0.07, *p* = 6.2 x _10_-13.

### 3. Pharmacological blockade of muscarinic ACh and D1/D2 DA receptors both reduce locomotion-evoked astrocyte Ca^2+^

Our results so far characterize the relationship between DA, ACh, and astrocyte Ca^2+^ under baseline conditions. These data suggest that both neuromodulatory systems are associated with increased striatal astrocyte Ca^2+^. Given this conclusion, we sought to investigate how striatal astrocyte activity is affected by pharmacological blockade of receptor subtypes most likely to underly this neuromodulator-astrocyte axis^44–47^. We blocked receptor activity for either muscarinic ACh receptors with scopolamine (2 mg/kg i.p.) or D1/D2 DA receptors with combined SCH 23390 (0.2 mg/kg i.p.) and haloperidol (5 mg/kg i.p.). We performed these receptor blockade experiments in mice bilaterally expressing virally introduced gfaABC1D-cyto-GCaMP6f and either GRAB-DA2m or GRAB-ACh3.0 (see **Fig 1a-c**). We then trained mice to locomote on a motorized wheel and recorded fiber photometry signals while varying locomotion durations either under receptor blockade or vehicle control conditions (**Fig 3a, d**). Under muscarinic receptor blockade with scopolamine, locomotion-elicited astrocyte events were reduced in comparison to saline control at all tested locomotion durations (**Fig 3b, c**). Under D1/D2 receptor blockade with SCH 23390 and haloperidol, astrocyte events were reduced only in the 10 and 20s conditions (**Fig 3e, f**), possibly due to the slower kinetics of DA-astrocyte correlation observed previously (**Fig 1i** and **Fig 2f**). Thus, systemic pharmacological blockade of both ACh and DA receptor activity compromises astrocyte Ca^2+^ activity evoked by locomotion, albeit to differing degrees.

**Figure 3:**
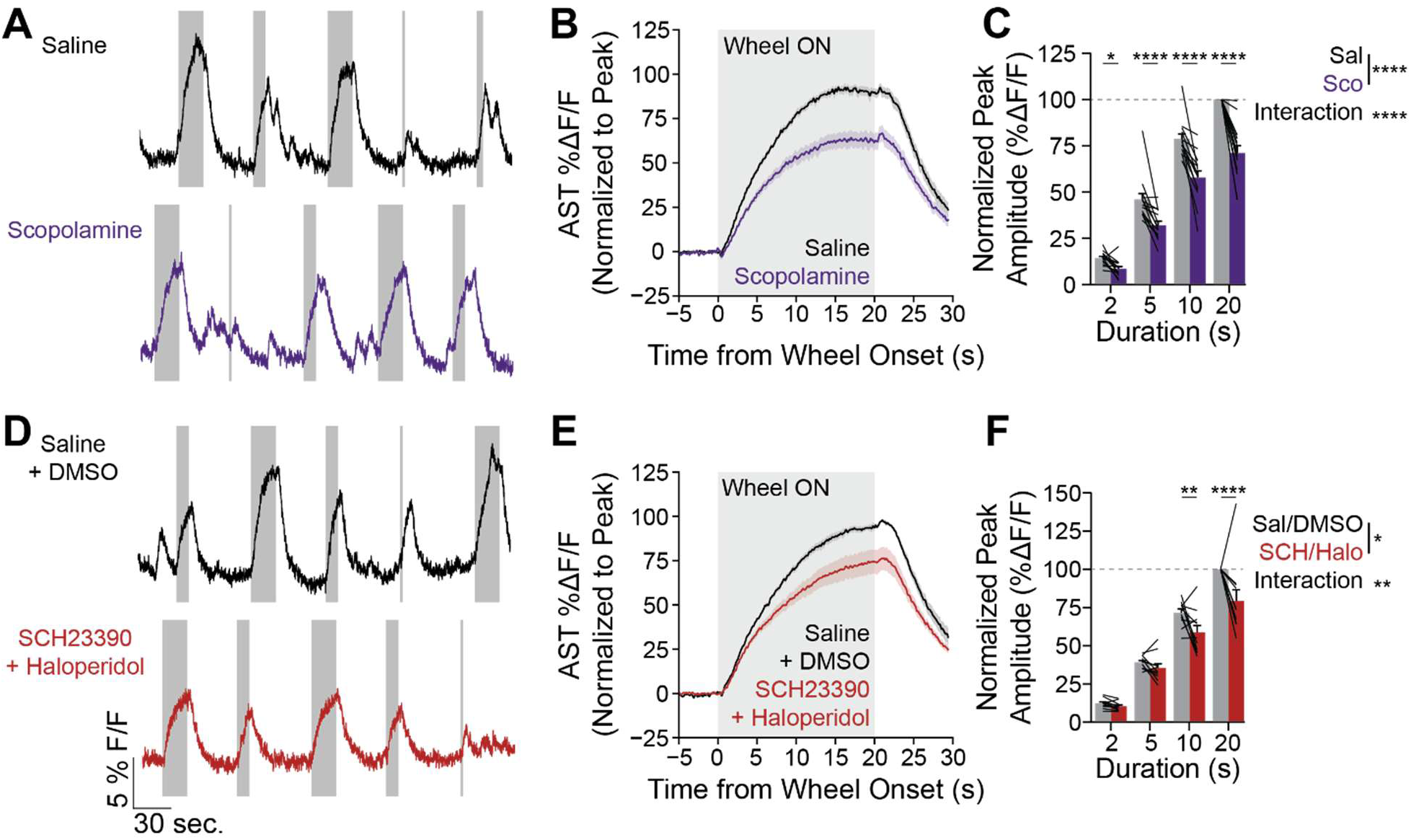
Pharmacological blockade of muscarinic and D1/D2 receptors reduces locomotion-evoked astrocyte Ca^2+^ activity. (**A**) Example traces of gfaABC1D-cyto-GCaMP6f activity under saline control (top) and muscarinic receptor antagonism with scopolamine (2 mg/kg i.p., bottom). Shaded portion represents motorized locomotion. (**B**) Animal averaged trace of astrocyte Ca^2+^ activity under saline or scopolamine injection for the 20s duration. Data shown are normalized to the animal averaged 20s saline ΔF/F max value. (**C**) Quantification of astrocyte Ca^2+^ reduction with muscarinic receptor antagonism (two-way ANOVA interaction *F*(3, 39) = 22.65, *p* = 1.2 × 10^−8^, main effect of scopolamine *F*(1, 13) = 40.82, *p* = 2.4 × 10^−5^. Sidak’s multiple comparisons test for saline vs scopolamine 2s *p* = 0.033, 5s *p* = 2.2 × 10^−7^, 10s *p* = 8.3 × 10^−11^, 20s *p* < 1.0 × 10^−15^, n = 11 mice). (**D**) Example traces of gfaABC_1_D-cyto-GCaMP6f activity under 2.5% DMSO in saline control (top) and D1/D2 antagonism with SCH 23390 (0.2 mg/kg i.p.) and haloperidol (5 mg/kg i.p., bottom). Shaded portion represents motorized locomotion. (**E**) Animal averaged trace of astrocyte Ca^2+^ activity under 2.5% DMSO in saline or SCH 23390/haloperidol injection for the 20s duration. Data shown are normalized to the animal averaged 20s 2.5% DMSO ΔF/F max value. (**F**) Quantification of astrocyte Ca^2+^ reduction with D1/D2 receptor antagonism (two-way ANOVA interaction *F*(3, 30) = 5.40, *p* = 0.0043, main effect of SCH 23390 + haloperidol *F*(1, 10) = 8.67, *p* = 0.014. Sidak’s multiple comparisons test for 2.5% DMSO vs SCH 23390 + haloperidol 2s *p* = 0.98, 5s *p* = 0.81, 10s *p* = 0.0077, 20s *p* = 2.1 × 10^−5^, n = 15 mice).

### 4. Closed loop optogenetic inhibition of striatal cholinergic interneurons reduces locomotion-evoked astrocyte Ca^2+^ activity

Our data suggest that high ACh release may trigger astrocyte Ca^2+^ increases, given its faster time to peak relative to dopamine release (**Fig 1i**). The primary source of ACh release in the DLS is from cholinergic interneurons (CINs) which make up ∼2-3% of the neuronal population. Given our results so far indicating a role for ACh activity in astrocyte Ca^2+^ elevations, we hypothesized that perturbing CIN activity may reduce astrocyte Ca^2+^. To test this possibility, we expressed the Cre-dependent red-shifted halorhodopsin hSyn-FLEX-Jaws-mCherry or control hSyn-DIO-mCherry in mice expressing Cre under the choline acetyltransferase promoter (ChAT-Cre) co-injected with gfaABC_1_D-cyto-GCaMP6f into DLS (**Fig 4a**). This experimental strategy specifically labeled striatal CINs with Jaws or mCherry constructs to allow for precise control of CIN activity with optogenetics during simultaneous astrocyte recording (**Fig 4b**). To target locomotion evoked ACh activity, we designed a closed-loop experimental paradigm under which locomotion data was processed in real-time to trigger continuous laser illumination at 589 nm when locomotion activity rose above a threshold (**Fig 4b**). Laser illumination was triggered on random 50% of trials while the remaining trials were used as laser “OFF” control. Closed-loop optogenetic inhibition of CIN activity acutely reduced astrocyte Ca^2+^ activity in Jaws expressing mice compared to OFF trials (**Fig 4c, d**). In mCherry expressing control mice, laser illumination did not significantly affect astrocyte Ca^2+^ activity during the photostimulation period (**Fig 4e**). We note that photostimulation in mCherry expressing mice elicited a positive going artifact which was not observed in Jaws expressing mice, suggesting that we are likely underestimating the effectiveness of CIN inhibition on reducing astrocyte Ca^2+^ activity. When compared directly to controls, laser induced ΔAUC was significantly lower in Jaws expressing mice than in mCherry control animals (**Fig 4f**). These data show that inhibition of CIN activity results in acute decreases of astrocyte Ca^2+^ activity, evidencing a direct involvement of ACh status on astrocyte activity. Overall, our results demonstrate a role for both DA and ACh in recruiting astrocyte Ca^2+^, with particular emphasis on the activity of striatal CINs as astrocyte activators.

**Figure 4:**
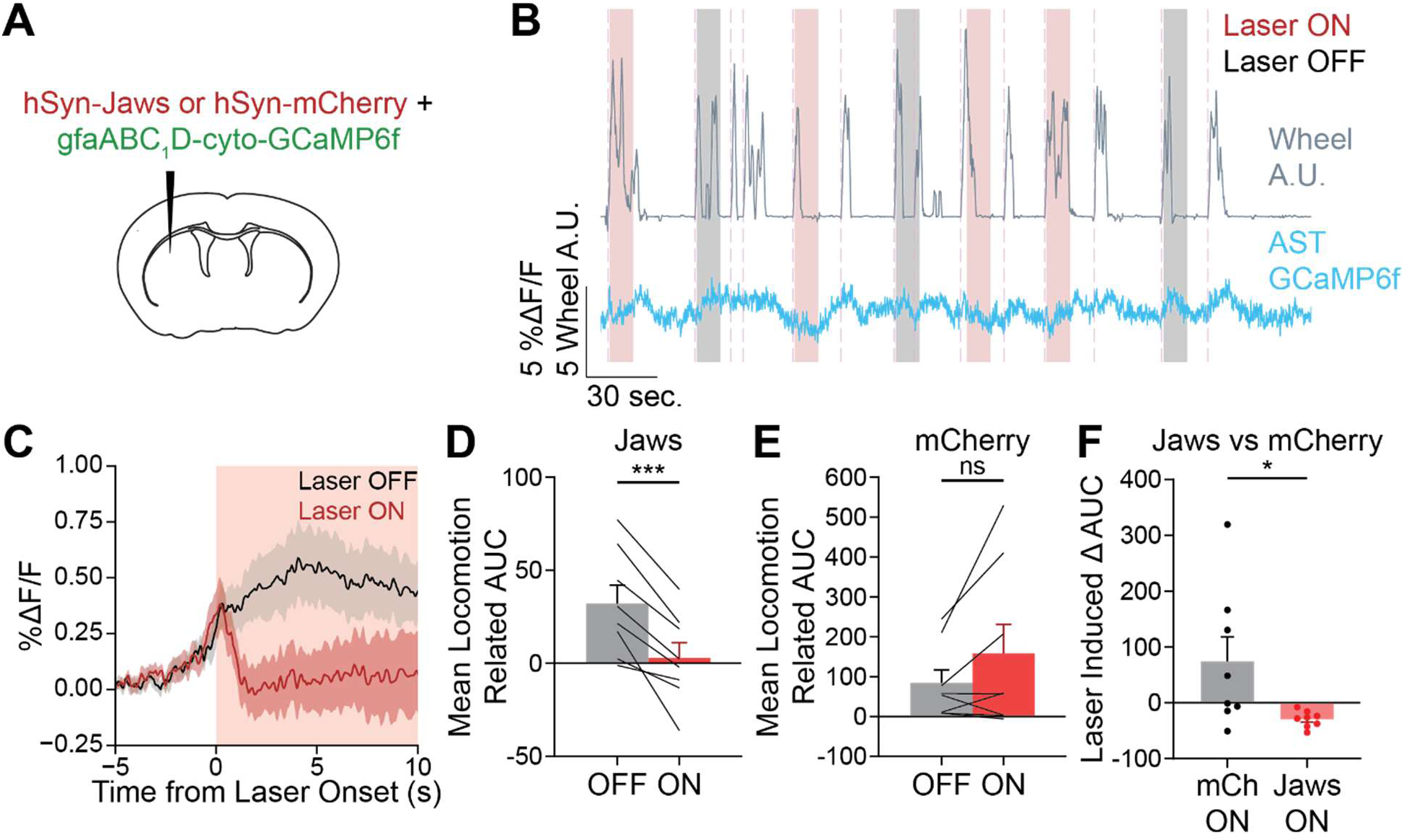
Closed-loop inhibition of CIN activity during locomotion acutely reduces astrocyte Ca^2+^ activity. (**A**) Experimental schematic showing viral construct co-injection strategy to express CIN-Jaws or CIN-mCherry alongside gfaABC_1_D-cyto-GCaMP6f. (**B**) Epifluorescence image showing Jaws expression in striatal CINs alongside astrocyte expression of GCaMP6f. (**C**) Example trace from a closed-loop inhibition experiment showing ON and OFF laser trials and locomotion detections. Note locomotion detections occurring without laser stimulation within the 30s lock out period post laser trial. (**D**) Astrocyte Ca^2+^ ΔF/F from CIN-Jaws expressing mice aligned to onset of laser trigger in ON and OFF trials showing acute negative deflection of astrocytes in ON trials. (**E**) Quantification of post-laser astrocyte Ca^2+^ ΔF/F AUC in ON and OFF trials from CIN-Jaws animals (paired t-test *p* = 0.00077, n = 8 mice). (**F**) Quantification of post-laser astrocyte Ca^2+^ ΔF/F AUC in ON and OFF trials from CIN-mCherry animals (paired t-test *p* = 0.14, n = 8 mice). (**G**) Comparison of laser-induced ΔAUC (ON – OFF) in CIN-Jaws and CIN-mCherry mice (Mann Whitney test *p* = 0.010, n = 8 CIN-Jaws and 8 CIN-mCherry mice).

### 5. DA denervation reduces locomotion evoked ACh and astrocyte Ca^2+^ activity

Based on our experimental evidence so far, we conclude that both ACh and DA are involved in driving locomotion-evoked increases in striatal astrocyte Ca^2+^. However, these experiments have only investigated the role of these neuromodulators in the healthy striatum. Whether this signaling axis is a potential pathological mechanism in states such as Parkinson’s disease remains an open question. To test whether and how this neuron-astrocyte communication axis is perturbed by pathological circuit conditions, we assessed the changes induced by DA denervation to ACh and astrocyte Ca^2+^ utilizing the 6-OHDA unilateral MFB lesion model^48^. Transgenic mice expressing GCaMP8s under control of the Aldh1L1 promoter were injected with rACh1.7 into both hemispheres and a bilateral cannula was placed above the injection sites. This strategy allowed us to record and compare within animal ACh release and astrocyte Ca^2+^ activity in affected and unaffected striatal hemispheres along with between-animal comparisons to sham injection controls (**Fig 5c**). We recorded from lesion animals during voluntary locomotion and aligned ACh and astrocyte Ca^2+^ activity to onset of locomotions. In this paradigm, we detected reduced astrocyte Ca^2+^ in the ipsilateral hemisphere and reduced ACh activity in both hemispheres when compared to sham treated animals (**Fig 5a, b**). We note that these data are difficult to interpret since locomotor activity from lesioned animals was expectedly lower when compared to sham treated animals (**Fig 5a**, bottom), necessitating the use of motorized locomotions of controlled durations to evoke comparable movement to sham (**Fig 5c**). In agreement with our previous work using a membraned-tagged version of GCaMP6f^23^, we found that astrocyte Ca^2+^ peaks from the ipsilateral hemisphere were decreased compared to sham and the contralateral hemisphere (**Fig 5d-f**). The ACh activity in the lesioned hemisphere showed a marked decrease when compared to both sham and control hemisphere (**Fig 5g-i**). Instead of a rapid rise in ACh release observed on the unlesioned side, ACh release in the lesioned hemisphere showed a negative dip right after locomotion onset and strongly reduced locomotion AUC in comparison to both sham and contralateral control (**Fig S2**). Thus, DA denervation leads to failure of ACh release to spike at locomotion and remain elevated during the locomotion bout. This, along with a lack of DA altogether, results in decreased subsequent astrocyte Ca^2+^ activity. Counteracting this ACh signaling deficit by directly augmenting cholinergic signaling in astrocytes improved locomotion-evoked Ca^2+^ activity and asymmetrical motor behavior characteristic of this animal model^23^. Together, our results show that the ACh-DA-astrocyte axis is a critical communication system for striatal circuit function and that the behavioral impact of lost neuromodulation may be restorable by directly targeting astrocyte Ca^2+^ activity.

**Figure 5:**
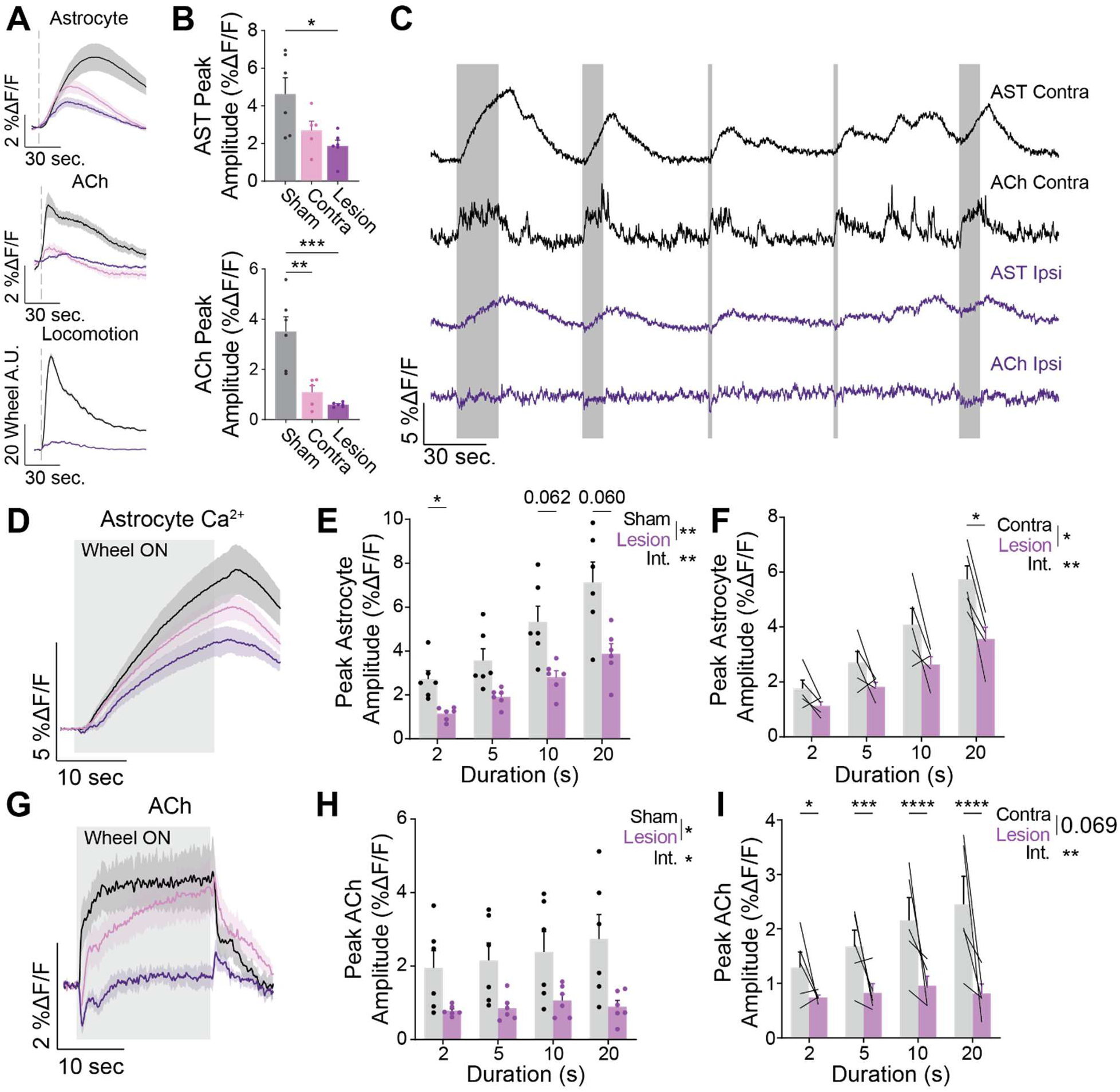
DA denervation causes dysfunctional locomotion related ACh release and reduced astrocyte Ca^2+^ activity. (**A**) Aligned animal average traces of astrocyte Ca^2+^ (top), ACh (middle), and locomotor (bottom) activity from sham treated and lesioned animals recorded from the ipsilateral and contralateral hemispheres. (**B**) Peak astrocyte ΔF/F values post-locomotion onset from sham, ipsilateral, and contralateral hemispheres (one-way ANOVA *F*(2, 14) = 5.565, *p* =0.0167; Tukey’s multiple comparisons test sham vs ipsilateral lesion *p* = 0.0148, sham vs contralateral lesion *p* =0.1103, contralateral vs. ipsilateral *p* = 0.6301) (top). Peak ACh ΔF/F values from sham, ipsilateral, and contralateral hemispheres (one-way ANOVA *F*(2, 14) = 16.77, p = 0.00019; Tukey’s multiple comparisons test sham vs ipsilateral p = 0.00026, sham vs contralateral p = 0.0021, contralateral vs ipsilateral p = 0.7768 (bottom). (**C**) Five minutes of example recording from a lesioned animal during motorized locomotions of varied duration (shaded areas represent locomotion epochs) showing ipsi- and contralateral ACh and astrocyte Ca^2+^ ΔF/F activity. (**D**) Aligned astrocyte Ca^2+^ ΔF/F activity during 20s motorized locomotion trials from sham, ipsilateral, and contralateral hemispheres. (**E**) Peak astrocyte Ca^2+^ ΔF/F values stratified by motorized locomotion duration from sham and ipsilateral lesion hemispheres (two-way ANOVA interaction *F*(3, 30) = 4.848, *p* = 0.0072; main effect of lesion status *F*(1, 10) = 11.45, *p* = 0.0070; Sidak’s multiple comparisons test 2s locomotion *p* = 0.0283, 5s *p* = 0.1022, 10s *p* = 0.0595, 20s *p* = 0.0616. (**F**) Peak astrocyte Ca^2+^ ΔF/F values from contralateral and ipsilateral striatal hemispheres from lesioned mice (two-way repeated measures ANOVA interaction *F*(2.011, 8.044) = 14.36, *p* = 0.0022; main effect of hemisphere *F*(1.000, 4.000) = 9.481, *p* = 0.0370; Sidak’s multiple comparisons test 2s locomotion *p* = 0.3363, 5s *p* = 0.3778, 10s *p* = 0.2411, 20s *p* = 0.0276). (**G**) Same as (**D**) for ACh release in the 20s locomotion condition. **(H**) Same as (**E**) comparing sham vs lesioned striatal ACh release (two-way ANOVA interaction *F*(3, 30) = 3.480, *p* = 0.0280; main effect of lesion status *F*(1, 10) = 6.533, *p* = 0.0286; Sidak’s multiple comparisons test 2s locomotion *p* = 0.2095, 5s *p* = 0.1513, 10s *p* = 0.2311, 20s *p* = 0.1488). **(I**) Same as (**F**) for ACh release from lesioned and contralateral hemispheres (two-way repeated measures ANOVA interaction *F*(3, 12) = 8.976, *p* = 0.0022; main effect of lesion status *F*(1, 4) = 6.112, *p* = 0.0688; Sidak’s multiple comparisons test for 2s locomotion *p* = 0.0161, 5s *p* = 0.00059, 10s *p* = 2.3 × 10^−5^, 20s *p* = 8.4 × 10^−7^).

## Discussion

There is a growing interest in neuronal-astrocyte communication mechanisms which has identified norepinephrine (NE) as a major driver of cortical astrocyte Ca^2+^ activity^38,39,41,42,49–51^. However, the striatum is known to be largely devoid of NE activity, something we recently confirmed with photometry recordings of NE release ^36,37^. Our present study helps to expand on this paradigm of neuronal signals interacting with local astrocyte Ca^2+^ activity into a region with it’s own specialized neuromodulatory system. In the absence of NE activity, the striatum is in a constant state of ACh and DA fluctuation which is often anticorrelated on fast, sub-second timescale^30,35,52^. Here, we show that ACh and DA display a distinct anticorrelated relationship directly after locomotion onset which is then followed by sustained period of positive correlation, which occur approximately during the rising phase of astrocyte Ca^2+^ activity (**Fig 1, 2d-f**). We determined that ACh and DA both affect the subsequent level of astrocyte Ca^2+^ activity which occurs during voluntary locomotions (**Fig 2g-m**) and that pharmacological blockade of candidate GPCR subtypes for ACh and DA both reduce locomotion-evoked astrocyte Ca^2+^ activity (**Fig 3**). Finally, we show that acute optogenetic inhibition of CIN activity reduces locomotion-evoked astrocyte Ca^2+^ responses, establishing a role for CIN activity’s direct relationship to subsequent astrocyte activation (**Fig 4**). These data together support a model for DLS astrocyte activity which is attuned to the striatal neuromodulatory millieu in a complex manner, with roles for both DA and ACh supported by our results.

We also demonstrate that this regional neuromodulator-astrocyte relationship is impaired when DA is removed from the circuit with unilateral 6-OHDA lesioning. We show that, when compared to the contralateral hemisphere and to sham treated controls, ACh activity in the DA denervated DLS fails to rise at locomotion onset (**Fig 5**). Given our results from **Fig 3** and **4**, it is reasonable to conclude that this failure of ACh to be engaged by locomotion results in the lowered astrocyte Ca^2+^ activity we have previously reported^23^ and replicated here (**Fig 5**). These data expand our understanding of the striatal neuron-astrocyte communication system and establish its role in PD pathology.

There is a growing body of evidence that an important consequence of neuromodulator actions in the CNS is signaling to local astrocytes^16,44,50,53^. Here, we interrogated the effects of the ACh-DA axis in the DLS in the context of astrocyte Ca^2+^ elevations, demonstrating that this signaling mechanism is attuned to the specific context of the DLS. Both ACh and DA peak levels predicted increased subsequent astrocyte Ca^2+^ levels (**Fig 2g-j**) and astrocytes displayed sensitivity to both muscarinic and dopaminergic receptor antagonism (**Fig 3**). Our prior work demonstrated that chemogenetic stimulation of muscarinic M4 signaling restores bilateral paw usage in 6-OHDA lesioned animals^23^. We interpret these results together as evidence of an underlying mechanism such that ACh and DA activity in the healthy striatum both signal for state-dependent astrocyte Ca^2+^. Given that our results also indicate dysfunction of this system (**Fig 5d-i**) and therapeutic action when normalized, we conclude that this system is critical for proper DLS circuit function. Underlying this model is the assumption that astrocyte Ca^2+^ elevations have some consequence on local SPN activity from which arises the benefit of chemogenetic M4 stimulation. Most likely, this would involve gliotransmitter release and receptor activation in SPNs^10,54^. It is also possible that distal actions consequent to astrocyte Ca^2+^ elevation are not dependent on neuronal mediation given that astrocytes form their own gap junction coupled network capable of inter-nucleus communication^55^. Further investigation is needed to elucidate the mechanisms downstream of astrocyte Ca^2+^ elevation which confer therapeutic benefit in DA denervation.

Our results in regard to ACh and DA recruitment of astrocyte Ca^2+^ can be interpreted to represent continuous astrocyte integration of both neuromodulators simultaneously. However, it is important to note that we employed fiber photometry to assess these systems *in vivo*, a technique which necessarily records bulk activity across a large area of tissue^56^. We cannot therefore resolve whether our co-imaging astrocyte responses, nor our pharmacological effects, are occuring at the same cell over time. Indeed, it is plausible that the effects demonstrated here represent two populations of astrocytes, likely in communication with each other locally, arising as decreases in overall bulk signal. Either possibility represents an intriguing line of future investigation.. Further, our results here suggest that even in the DA denervated striatum, there is residual astrocyte Ca^2+^ during movement (**Fig 5d-f**), indicating that ACh and DA are not solely responsible for astrocyte recruitment. What factors are still engaged in this context is not resolvable from our current study, but there is evidence for GABAergic communication from SPNs to striatal astrocytes^9^. Other possible factors which result in DLS astrocyte Ca^2+^ include glutamatergic inputs from cortex and thalamus^25^. Characterizing these signaling axes represent other directions of future work.

Astrocytes in the context of NE activity have been shown to “gate” Ca^2+^ increases elicited by other neurotransmitters^38^. In our study, we do not directly assay whether a mechanism such as this is present in striatal astrocytes. However, viewed in the lens of astrocyte gating, our results suggest that ACh may play a similar role in the striatum to NE in other regions. The first signal to rise among ACh, DA, and astrocyte Ca^2+^ is ACh, exhibiting a fast, transient spike before settling into an elevated tonic phase. This spike precedes both the peak of astrocyte Ca^2+^ and DA activity. It is possible that this early spike represents a ACh gating mechanism, sensitizing astrocyte activity to the following peak in DA activity. This hypothesis would explain the temporal relationship of our investigated signals as requiring fast DA decreases to allow ACh to spike, signaling to the local astrocyte population to sensitize to the slower increase in DA. This possibility requires further data to fully evidence as fiber photometry’s bulk signals do not provide the resolution necessary to assess the single-cell reaction to neurotransmitter exposure. An alternative interpretation of our results is that individual astrocytes directly sample both ACh and DA activity, with the elevated post-locomotion level of both neuromodulators engaging astrocyte Ca^2+^ increases simultaneously. In this interpretation, the loss of DA with 6-OHDA and the loss of ACh activity at locomotion onset cooperatively decrease subsequent astrocyte levels. Indeed, our pharmacological experiments blocking receptor activity for either ACh or DA do not decrease astrocyte Ca^2+^ levels to the same degree as 6-OHDA lesioning (**Fig 3c, f** vs **Fig 5e, f**).

Astrocytes are known to be important for state switching of CNS circuitry^19,57^. For example, astrocyte Ca^2+^ activity increases in cortical Up-Down state transitions^58,59^, sleep stage transitions^60,61^, and slow wave oscillations^62^. Akinesia is a major symptom of PD in which patients exhibit an inability to initiate movements and is associated with aberrant neuronal state dynamics^63,64^. In rats, 6-OHDA lesioning causes circuit deficits in areas innervated by SPNs such that beta activity increases and gamma activity decreases which is associated with the onset of motor symptoms post-lesion^65^. With this context, our data may be interpreted as a possible state-switching mechanism which engages astrocyte activity. The onset of locomotion induces a rapid increase of ACh release and decrease of DA release (**Fig 1f-i** and **Fig 2e**), followed by a steady increase in astrocyte Ca^2+^ which lasts until the cessation of locomotion. It is possible that this pattern represents a possible state transition motif associated with the to switch into a locomotor phase where proper integration of cortical, thalamic, and SNpc DA information can occur. Astrocyte Ca^2+^ in this interpretation would be a consequential and slower signal which responds to this neuromodulatory pattern. The ACh portion of the motif becomes weaker or even non-existent with DA lesioning (**Fig 5a, b** and **g-i**). It is reasonable therefore to conceptualize the neuron-astrocyte communication in the DLS during locomotion as a state transition to initiate locomotion which is dysfunctional in Parkinsonian conditions, possibly contributing to akinetic motor symptoms.

Striatal CIN activity remains an open area of investigation in PD. The strongest effect observed in DA lesion experiments has been increased frequency of CIN action potentials post-lesioning^33,35^. This is often described as a state of hypercholinergic activity post-lesion, but there is an emerging view of this phenomenon as resulting in decreased ACh-mediated M4 activity^66,67^. Our results show a failure of ACh increasing at locomotion onset (**Fig 5g-i**) which does not directly address whether chronic levels of ACh are elevated or depressed. Instead, our results evidence a model where ACh activity appears discoordinated. Without DA activity (the dip of which is the earliest event post-locomotion of our studied signal features, **Fig 1i**) there is a failure of ACh activity to spike. One explanation of this phenomena is that the DA dip is necessary to remove CIN D2R inhibition to allow for the appropriate spiking of local ACh levels. Without DA signaling, CIN spike frequency increases may evidence a less synchronized environment which results in lessened astrocyte Ca^2+^ levels. Taken together, our results here expand the view on ACh activity in the lesioned striatum to include a reduced astrocyte Ca^2+^ status as a possible pathologic outcome.

Beyond the future directions mentioned above, there are several directions of investigation worthy of note. Firstly, characterization of the signaling pathways engaged by both ACh and DA in astrocytes may elucidate targets for therapy which engage native signaling systems. Astrocyte cAMP signaling has been demonstrated to play important roles in memory formation and is involved in neurodegenerative pathologies^68,69^. Future work should investigate genetic expression and signaling changes which result as a consequence of DA loss. Another future direction suggested by our results is investigating the nature of ACh dysfunction we observed (**Fig 5g-i**). We show that ACh activity in the DA denervated striatum is strongly affected compared to intact sham controls, showing that DA is crucial for proper ACh signaling in the DLS. While sham animals display a sharp post-movement onset peak in ACh release, 6-OHDA lesioned animals instead exhibit a dip in activity at the same time. From there, ACh levels fail increase above the pre-movement baseline. What causes this dysfunction represents an intriguing direction for understanding striatal circuitry and may reveal important findings in regards to how Parkinsonian deficits arise in the DLS circuitry. Therapeutic avenues to restore proper ACh activity may represent an important, non-dopaminergic intervention. The frontline treatment for PD is DA replacement with the precursor levodopa (L-DOPA); however, continued use carries the risk of L-DOPA induced dyskinesia^70–72^. Treatments which do not confer this side-effect are therefore necessary for improvement of patient quality of life.

Overall, our study indicates that striatal astrocytes are engaged by both ACh and DA release, expanding the view of neuromodulator-astrocyte communication into a unique neuromodulatory environment. Our results provide insight into astrocyte regulation of DLS circuitry and evidence a neuron-astrocyte communication system which becomes dysfunctional in disease pathology. These results have important implications for understanding striatal function and for treatment of PD.

## Supplemental Figures

**Supplemental Figure 1:**
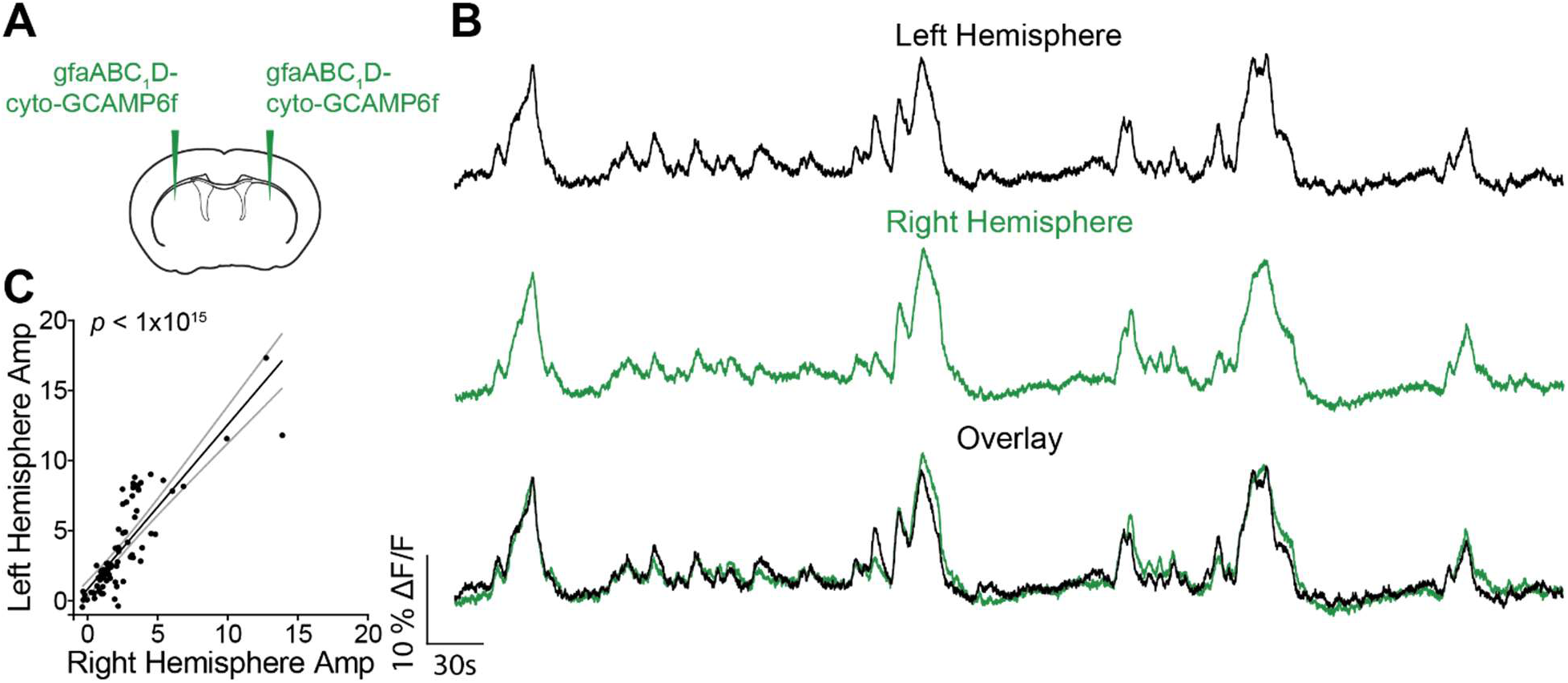
Bilateral astrocyte Ca^2+^ activity is highly correlated. (**A**) Experimental schematic for expression of bilateral gfaABC1D-cyto-GCaMP6f for interhemispheric recording of astrocyte activity. (**B**) Example traces from the left (top) and right (middle) DLS hemispheres and overlay of both (bottom). (**C**) Detected event amplitudes across hemispheres and linear regression best fit line (linear regression *r*^2^ = 0.72, *p* < 1 × 10^−15^).

**Supplemental Figure 2:**
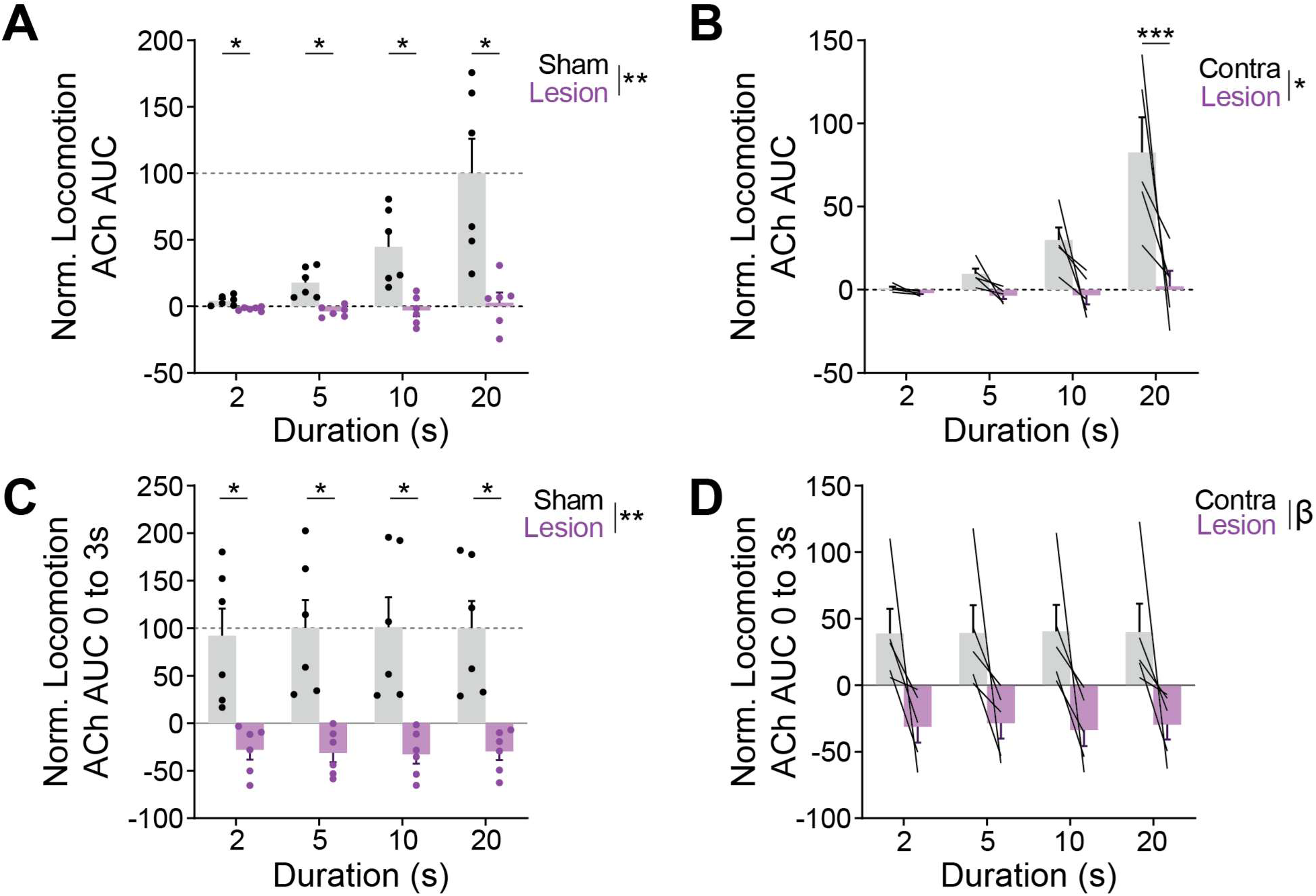
ACh release during locomotion and at movement onset show inhibition in DA denervation. (**A**) ACh release AUC during motorized locomotion normalized to mean 20s sham AUC (two-way ANOVA interaction *F*(3, 30) = 12.15, *p* = 2.2 × 10^−5^, main effect of lesion *F*(1, 10) = 14.35, *p* = 0.0036; Sidak’s multiple comparisons test for 2s duration *p* = 0.033, 5s *p* = 0.016, 10s *p* = 0.031, 20s *p* = 0.047). (**B**) ACh release during locomotion from lesion contra-and ipsilateral hemispheres normalized to mean 20s sham AUC (two-way repeated measures ANOVA interaction *F*(3, 12) = 7.69, *p* = 0.0039, main effect of hemisphere *F*(1, 4) = 8.17, *p* = 0.046; Sidak’s multiple comparisons test 2s duration *p =* 0.99, 5s *p* = 0.78, 10s *p* = 0.078, 20s *p* = 0.00012). (**C**) ACh release AUC during the first 3s post locomotion normalized to 20s sham AUC from 0 to 3s (two-way ANOVA interaction *F*(3, 30) = 1.03, *p* = 0.3928, main effect of lesion *F*(1, 10) = 17.61, *p* = 0.0018; Sidak’s multiple comparisons test for 2s duration *p* = 0.028, 5s *p* = 0.020, 10s *p* = 0.027, 20s *p* = 0.020). (**D**) ACh release AUC during the first 3s after locomotion onset from lesion contra- and ipsilateral hemispheres normalized to 20s AUC (two-way repeated measures ANOVA *F*(3, 12) = 0.57, *p* = 0.64, main effect of hemisphere *F*(1, 4) = 6.17, *p* = 0.068.

## Methods

All experiments and procedures performed using animals were approved by the Rutgers University Institutional Animal Care and Use Committee (Protocol #202000004). All mice in the study were bred in-house from progenitors ordered from the Jackson Laboratory. Strains used in this study were C57BL/6J (JAX 000664), ChAT-cre (JAX 031661), GCaMP8s (JAX 037952), and Aldh1l1-cre/ERT2 (JAX 031008) back-crossed onto a 6J background as confirmed by genetic monitoring (Transnetyx). In some experiments, astrocytes were labeled using a cross of GCaMP8s mice with Aldh1l1-cre/ERT2 mice and recombinase activation with 5 days of tamoxifen induction. Following surgery, mice were housed under a reverse light-dark cycle (8am-8pm) in standardized cages and given food and water *ad libitum*.

### General Stereotaxic Surgery Procedures

Procedures were conducted in accordance with our prior work^23,73^. Anesthesia was induced under 4% isoflurane and maintained at 1%-2% isoflurane and adjusted as needed for surgery. Mice were then mounted on a stereotaxic frame warmed to ∼37°C (Stoelting). Anesthesia was continued Analgesia was provided for 72 hours using extended-release buprenorphine (Ethiqa XR, 3.25 mg/kg). Bupivacaine (1 mg/kg) was given subcutaneously under the scalp. The scalp was then depilated using Nair and disinfected using alternated swabs of betadine and 70% ethanol. The scalp was removed and the skull was leveled relative to bregma and lambda. Mice were kept warm while recovering from anesthesia and were monitored during recovery for at least 3 days post-surgery.

### 6-OHDA Infusion

At least one week prior to surgery, mice were presented with additional dietary supplementation consisting of Pediasure, peanut butter, DietGel, and HydroGel to combat neophobia and increase body weight. Mice were injected intraperitoneally with desipramine hydrochloride (HCl) and pargyline HCl (10 mL/kg) in 0.9% sterile saline 30 min before 6-OHDA infusion^48^. A burr hole was drilled in the skull and 6-OHDA HBr (3.0mg/mL) in 0.02% ascorbic acid/0.9% sterile saline solution was pressure injected (Nanoject III, Drummond) into the MFB at the following coordinates relative to bregma: AP: −1.2, ML: +1.2, DV: −5.0 mm. For sham surgeries, animals received both the I.P. injection of desipramine and pargyline and a pressure injection of the vehicle solution into the MFB. Injections were done over the course of 10 minutes with the needle being left in place for another 10 minutes after. Additional injections and implants occurred during the same surgery depending on the experiment (see below). Lesion mice were monitored for at least 14 days post-surgery. Mice were provided with a heating pad under half of the home cage to assist in maintaining body temperature. Mice were presented with Pediasure using a pipette and given subcutaneous injections of 2.5% dextrose in 0.9% sterile saline or lactated Ringer’s solution. Additional dietary supplementation consisting of peanut butter, DietGel, and HydroGel was placed in the cage daily for the life of the animal.

### AAV Injections

A burr hole was made above the DLS using a handheld drill. Viral constructs were pressure injected (Nanoject III, Drummond) into the DLS using the coordinates AP: 0.4, ML: 2.3, DV: −2.8 relative to bregma. We utilized the following constructs according to experimental designs (see **Results**): gfaABC_1_D-cyto-GCaMP6f (Addgene, 52925-AAV5), hSyn-GRAB-ACh3.0 (Addgene, 121922-AAV9), hSyn-GRAB-DA2m (Addgene, 140553-AAV9), hSyn-GRAB-rACh1.7 (BrainVTA, PT-5488), hSyn-GRAB-rDA1m (Addgene, 140556-AAV9), and CAG-FLEX-rc [Jaws-KCG-tdTomato-ER2] (Addgene, 84446-AAV5). Following pressure injection, the pipette was kept in place for 5 minutes before being slowly withdrawn.

### Fiber Optic Implants

Following AAV injection into the DLS, a bilateral fiber optic cannula (Ø1.25 mm ceramic ferrule, 400 μm Core, 0.50 NA5.0 pitch dual site cannula, Doric Lenses) was fixed onto a stereotaxic cannula holder (ThorLabs) and slowly lowered into the brain at the same coordinates as the viral injection. For unilateral fiber experiments (**Fig 4**), a single fiber of the same specifications was implanted (RWD Life Sciences). The fiber was then implanted 0.1 mm above the AAV injection, and dental cement (MetaBond, Parkell) mixed with mica powder was used to secure the cannula in place. A custom headplate (eMachineShop) was placed on the skull using cyanoacrylate (Loctite) and fixed in place using dental cement (MetaBond, Parkell) to allow for head-fixed experiments. A subcutaneous injection of 0.9% sterile saline was administered, and mice were kept warm on a heating pad while recovering from anesthesia.

### Fiber Photometry

Mice were given two weeks post-surgery to allow for surgical recovery and viral expression. Photometry signals were recorded using Synapse software on a Tucker-Davis Technologies RZ10x. Green fluorophores were excited at 465 nm alongside isosbestic excitation at 405 nm while red fluorophores were recorded at 560 nm. A Doric minicube was used to collate excitation wavelengths and to separate emission wavelengths into separate sensors. A patch cord was attached to the implanted ferrule using a ceramic coupling sleeve (Doric Lenses). Signals were then analyzed using custom Python scripts based on previously established methods^74,75^. The emission and excitation signals were downsampled to ∼100Hz. The isosbestic channel was fitted to the green sensor-dependent signals using least-squares polynomial fitting of order 1 and red sensor activity was corrected using a fitted exponential. Signal ΔF/F was calculated as ΔF/F = ((Signal – Control) / Control) × 100.

### Optogenetic Experiments

Closed loop optogenetic control was achieved using a circular wheel attached to a rotary encoder (see below) with live Arduino wheel movement data used to trigger laser illumination. A custom MATLAB script monitored incoming wheel movement data and triggered Arduino controlled laser onset. Jaws was activated with a Shanghai Lasers 589 nm orange laser with a continuous 15 mW pulse lasting 10s. Identity of trials was determined probabilistically at locomotion detection to either trigger laser illumination or not, analyzed as “ON” and “OFF” trials. After trial completion, a 30s timeout period was enforced until the next could be triggered, and full sessions lasted 40 minutes.

### Voluntary Head-Fixed Locomotion

Voluntary locomotion was conducted as previously described^23^. In brief, mice were head-fixed on a custom-made wheel attached to a rotary encoder (YUMO) placed at a 20° angle to record locomotor activity. The rotary encoder was attached to an Arduino Uno running a custom sketch and experiments were run using a custom MATLAB script. Data were synchronized by sending TTLs from the Arduino to the RZ10X during the recording. Animals were habituated to the voluntary wheel for a minimum of three sessions until they consistently ran forward, and mice that failed to habituate were excluded from analysis. Individual locomotion bouts were identified by finding continuous wheel speed samples that were above a threshold (2 A.U.) for a minimum duration of 2 s and were preceded by 5 s of wheel quiescence.

### Motorized Wheel Head-Fixed Locomotion

Motorized locomotion was conducted using the same setup as previously described^23^. In brief, mice were head-fixed on a custom-made wheel as described above for voluntary locomotion with a D/C motor in place of the rotary encoder. Motorized locomotion trials occurred for a randomized duration of 2s, 5s, 10s, or 20s with an inter-trial interval of one minute. Ten trials were conducted for each of the above lengths over the course of each session. Animals were habituated as described previously. For pharmacology experiments the agents were injected 30-40 minutes before the session.

### Immunohistochemistry

Mice were perfused using 4% paraformaldehyde (PFA). Brains were extracted and kept in 4% PFA overnight before being moved to PBS-Azide the next day. Coronal slices of the dorsal striatum and substantia nigra were taken at a thickness of 60μm using a Compresstome vibratome (VF-310-0Z, Precisionary Instruments). Using 12-well plates, slices were first washed in PBS thrice for 15 minutes each before being incubated in 50% methanol in PBS for 30 min. Slices were then incubated in 1% hydrogen peroxide in PBS for 15 min before again being washed in PBS thrice for 15 min each. Tissue was then washed in 0.5% Triton-X in PBS for 30 min and blocked in a solution of 10% Normal Goat Serum with 0.2% Triton-X in PBS for 90 min. For the primary antibody, tissues were placed in the above blocking solution with chicken anti-TH 1:1000 (TYH-0020, Aves Labs) and left overnight on an orbital shaker at 4°C. The following day, slices were washed in PBS four times for 10 min each before being placed in a secondary of AlexaFluor 647 goat anti-chicken IgY 1:500 in PBS for two hours. Tissue was again washed in PBS thrice for 10 min each. Finally, using DAPI mountant, tissue was mounted on permafrost slides and covered with 24 x 55 mm No. 1 coverslips and stored at 4°C until imaging.

## Author Contributions

W. E. and R. H. designed experiments. W. E. performed and analyzed fiber photometry and histology experiments. R. H. analyzed fiber photometry experiments. C. J. and H. W. aided with fiber photometry experiments. A. V. and M. S. aided with histology experiments. W. Evans, H. W., and R. H. wrote the manuscript with edits from all authors. R. H. supervised the study and secured funding.

## Acknowledgements

This work was supported by the Rutgers IRACDA-INSPIRE Training Program (K12GM093854, W.E.), National Institute on Alcohol Abuse and Alcoholism (5R01AA030594-02 to R.H.), and New Jersey Health Foundation (R.H.). We thank the members of the Huda Laboratory for their feedback on this work. We thank the School of Arts and Science-Human Genetics Institute of NJ (SAS-HGINJ) Imaging Core Facility for use of their epifluorescence and confocal microscopes. We thank Molly Saunders Lucy for help with animal care and histological support.

## Data Availability Statement

All data and code generated as part of this study are available from the corresponding author upon reasonable request.

